# Graphical factorial surveys reveal the acceptability of wildlife observation at protected areas

**DOI:** 10.1101/580688

**Authors:** Jacopo Cerri, Elena Martinelli, Sandro Bertolino

**Affiliations:** Istituto Superiore per la Protezione e la Ricerca Ambientale, Piazzale dei Marmi, 57123, Livorno, Italy.; Dipartimento di Scienze della Vita e Biologia dei Sistemi, Università degli Studi di Torino, Via Accademia Albertina, 13, 10123, Torino, Italy

**Keywords:** factorial surveys, vignettes, acceptability, ibex, visitors, parks

## Abstract

Approaching large ungulates at protected areas is dangerous both for visitors and the animals. Nudging interventions can mitigate this issue, but conservationists need to be sure about which factors influence the acceptability of human-wildlife encounters, to design them.

In summer 2018, we recruited a sample of visitors (n = 205) at the Gran Paradiso National Park (Italy). They evaluated the acceptability of 9 digitally modified pictures, depicting a group of visitors observing an alpine ibex (*Capra ibex*) close to a trail. Pictures were characterized in terms of group size and distance from the ibex.

Observing ibexes was deemed to be acceptable if visitors were further than 25 meters from animals and when groups included less than 3 people. Approaching ibexes at 5 meters was always deemed to be unacceptable. Potential for Conflict Index (PCI) was constant across distance classes and it was generally low.

Our findings indicate that visitors share normative beliefs about the optimal distance and group size, that should characterize encounters with large ungulates, when visiting the park. These normative beliefs are crystallized, because previous encounters with ibexes did not affect the evaluation of each scenario and because the PCI was constant and low. We believe that behavioral interventions aimed at promoting respectful and safe human-ibex interactions can be enforced in areas where this interaction is critical, mostly in form of panels on hiking trails introducing normative pressures on visitors and motivating them to comply with rules.

## INTRODUCTION

Large mammals are iconic wildlife species that constitute a major attraction for tourists worldwide (Grünewald, Schleuning & Böhning-Gaese, 2016; Lindsey, Alexander, Mills, Romañach, Woodroffe, 2007; Penteriani et al., 2017b). Many people visit protected areas all around the world in the attempt to observe them in their natural habitat, for the most diverse reasons (Curtin, 2009, 2010). Unfortunately, observations are sometimes intrusive for animals and dangerous for people. Even without considering carnivores (e.g. bears Penteriani et al., 2017a) it is worth noticing that about 4 people got injured every year in the attempt to approach bisons (*Bison bison*) at the Yellowstone National Park (Miller & Freimund, 2017). On the other hand, when tourists number increase their continuous disturbance to wildlife could directly impact upon the well-being of the focal population, with disruption of daily behaviour including feeding, breeding, and resting (Holmes, Knight, Stegall & Craig, 1993; Geffroy, Samia, Bessa & Blumstein, 2015).

Negative interactions can be minimized in part by building adequate observation structures, adopting limiting regulations, or by fencing hiking trails. However, these solutions are hard to implement, and expensive to maintain, at the landscape scale. Their feasibility is limited to critical hotspots and their effectiveness strongly relies on visitors compliance with appropriate codes of conduct, which are can be violated.

In the last few years, nudging became pervasive in policy making and governance. Nudging relies on advances in the psychology, and it aims to guide human behavior maximizing collective well-being. It often does so through behavioral interventions exploiting cognitive bias or introducing social pressures through normative messages (Halpern, 2016; Kinzig et al., 2013; Thaler & Sunstein, 2009). Not surprisingly, nowadays protected areas regard nudging with growing interest, as it promises to make visitor behavior less impacting for the environment. For example, some parks tailor messages to particular segments of visitors (Miller, Freimund, Metcalf & Nickerson, 2018) or even experiment the effectiveness of communication on visitor behavior directly (Cialdini, 2003; Cialdini et al., 2006; Hockett, Marion & Leung, 2017). As our understanding of human behavior and influence strategies is steadily growing, there is little doubt that nudging will also become important for optimizing the management of protected areas and to achieve conservation goals. However, conservationists should not be too naive: poorly designed behavioral interventions can seriously backfire, resulting in a rapid escalation of deviant and undesirable behavior that worsen the initial situation (Schultz, Nolan, Cialdini, Goldstein, & Griskevicius, 2018). Behavioral interventions should always be designed after an informative phase. Developing easy-to-use and reliable tools to understand how and why people behave in a certain way is crucial at this stage.

The alpine ibex (*Capra ibex*) is an iconic ungulate characterizing many protected areas in the Alps. Being one of the largest mammal in Europe, highly visible living in open areas, and often confident caused by having been protected for more than one century, makes the species prone to being approached by hikers. This is particularly common in the Gran Paradiso National Park (Italy), the only area where the species survived at the beginning of the last century and source of all the populations in the Alps. Here the species reach high densities and is particularly confidence toward humans Visitors approach ibexes to observe them and to take pictures, as the species is the symbol of the park. The park is therefore a good place where to study humans and wildlife interactions. Various studies pointed out that visitors acceptability of human wildlife interactions is determined by crowding at observation sites and by distances between humans and wildlife (Anderson et al., 2010; Miller & Freimund, 2018; Prakash et al., 2018; Verbos et al., 2017). This study aims to apply a visual-based factorial survey (Anderson, Manning, Valliere, & Hallo, 2010; Miller & Freimund, 2018) to elicit the acceptability of these interactions, in a European context. We will show how a common understanding of acceptable human-wildlife can be elicited, to guide the implementation of panels targeting visitors.

## MATERIALS AND METHODS

We designed a factorial survey experiment (Auspurg & Hinz, 2014), where each respondent evaluated 9 pictures representing hypothetical situations where visitors observed an adult male alpine ibex. Ibex present strong sexual dimorphism, with males characterized by large body mass and long backwards-curving horns, reduced in females (Bergeron, Festa-Bianchet, von Hardenberg, & Bassano, 2008). Scenarios varied in the number of visitors (1, 3 and 6 people) and in the distance between visitors and the ibex (approximately 5, 25 and 50 meters). For each vignette, a trained enumerator asked respondents to rate the acceptability of the human wildlife interaction on a 5-points bipolar ordered scale with a neutral point, ranging from “Totally unacceptable” to “Totally acceptable”.

Factorial surveys are usually narrative, with each vignette describing a situation in a text. However, factorial surveys based on digital pictures can provide an extremely vivid and salient description of a hypothetical situation, especially when they depict something that can be found during a real experience. This maximizes the plausibility of a counterfactual situation, reducing cognitive burden and improving the overall quality of vignette evaluation (Auspurg & Hinz, 2014). In our case, we photographed a portion of a trail where alpine ibex can be observed, using a boulder as a reference point. A collaborator was asked to change its distance from the boulder from 5 meters, to about 25 and 50 meters. Then, we added a male alpine ibex picture and we also varied the number of people in the group observing the ibex (Fig. 1). Considered that our factorial design included 9 combinations only, respondents evaluated the entire range of potential scenarios. Pictures representing hypothetical scenarios were displayed in a random order, to control for ordering effects. The factorial survey included a final section collecting baseline details of respondents (e.g. age, level of education, sex, residence), as well as information about their frequency of park utilization, their previous experience with alpine ibexes, their recreational preferences (e.g. angling, hunting, photography) and their belonging to environmental NGOs. In July-August 2018, we intercepted a sample of visitors in three areas highly frequented by tourists in the Gran Paradiso National Park (Aosta, Italy; Fig. 2). Visitors were recruited on a voluntary basis and no reward was provided. They took approximately 5-10 minutes to complete the questionnaire and to evaluate all the vignettes. A complete version of the factorial survey, altogether with the pictures is available in the Supplementary Information (S1).

**Fig. 1.**
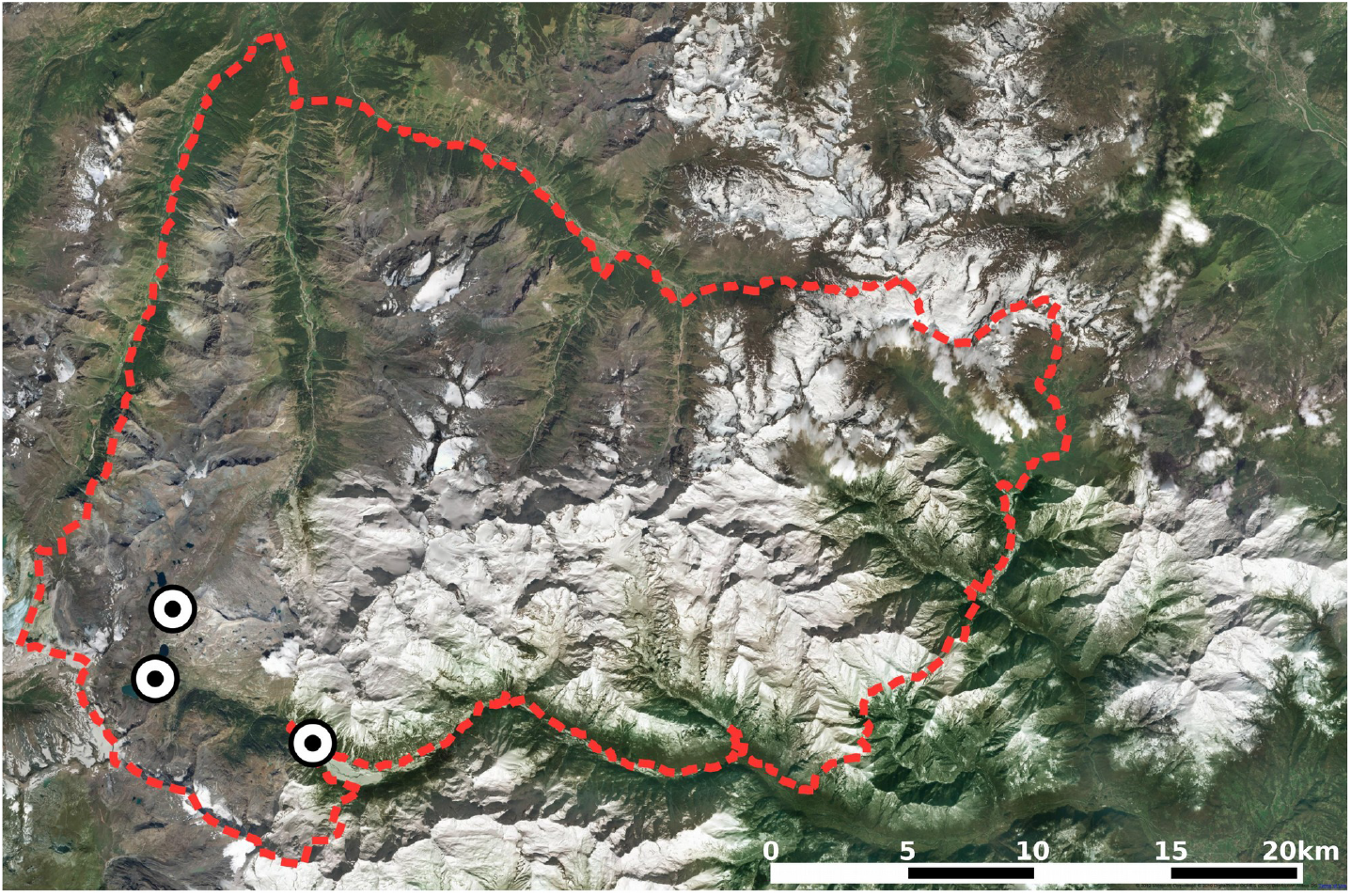
Map of the study area: boundaries of the Gran Paradiso National Park (dashed line) and locations where questionnaires were administered.

**Fig. 2.**
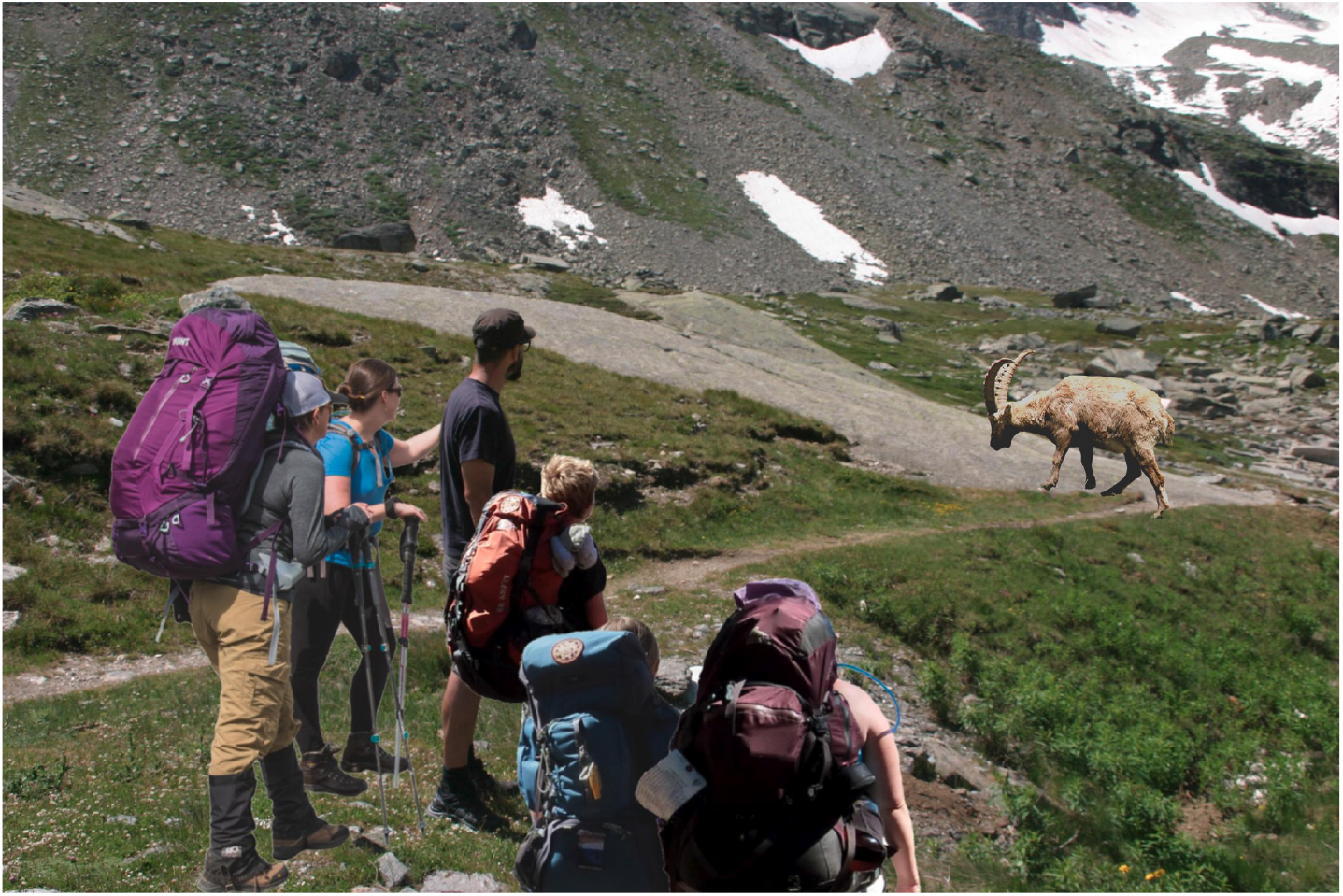
Example of a vignette, depicting o group of 6 visitors approaching an alpine ibex at 25 meters.

We used Bayesian Generalized Linear Mixed Modeling (GLMM) with a random intercept and random slopes for each predictor, to model how various predictors affected the perceived acceptability of each scenario. Predictors included the distance between visitors and the ibex, group size, previous experience with ibexes and a dummy variable measuring whether visitors practiced naturalistic photography or not. We included this latter predictor, because naturalistic photographers are a category that usually considers the issue of approaching animals in-depth and they can hold different beliefs from common visitors. Only 2% of respondents practiced recreational angling, there were no recreational hunters and 1.99% of visitors were members of environmental NGOs. Therefore, these variables could not be considered in the statistical analysis. An interaction term between being a photogapher, the distance from the ibex and group size was used. Our response variable measuring acceptability was measured on an ordered scale: there is no universal consensus on whether these scales should be treated as normally distributed or not (Bürkner & Vuorre, 2018; Liddel & Krushcke, 2017; Norman, 2010). To overcome this controversy we compared two concurring models, one based on a Gaussian distribution of the errors and one adopting a logit link function for ordinal responses. Model selection was based on leave-one-out cross validation and the widely applicable Bayesian Information Criterion (WAIC). Comparing these two models also enabled us to obtain insights about the overall fitness of the model, as some metrics like the R^2^ or the Intraclass Correlation Criterion (ICC) can be computed for Gaussian mixed models only.

We also used the Potential for Conflict Index (PCI2; Vaske, 2018) to assess the level of overall agreement about the various combinations of distances and group sizes. The PCI ranges between 0, when there is total consensus and all the answers have the same value, and 1, when the answers are equally distributed between two opposite values of the scale. We deemed the PCI to be informative about the presence of small groups of respondents who disagreed on some particular answers.

Statistical analyses were implemented on the software R and Bayesian multilevel modeling on the STAN software, through the ‘brms’ package (Bürkner, 2017). GLMMs had 4 MCMC chains running in parallel with 5000 iterations each. A complete R script is available in S2.

## RESULTS

We recruited a total of 205 visitors, who evaluated 1723 scenarios. Completion rate was rather high (94.31%). Respondents’ age was 37.20 ± 11.76 years (mean ± standard deviation), 51.41% of respondents were women and average education was high (8.80% of respondents had a secondary degree, 50.89% a high school diploma and 40.29% a university degree). 96.55% of respondents were visiting the park with a peer and 28.19% of them were visiting the park for the first time. Almost half of our sample (48.93%) had never seen an ibex at the park, before.

The GLMM with a binomial error strongly outperformed the Gaussian one, in terms of WAIC and leave-one-out cross validation. It was also clearly superior, when we compared the distribution of the observed outcomes with kernel density estimates of each replication from the posterior predictive distribution (Fig. 3). However, both the Gaussian and the binomial GLMM selected an identical final set of predictors, which included the distance between visitors and the ibex, group size, their interaction and the dummy variable classifying respondents as photographers or not. While the R^2^ and the ICC could not be computed for the final binomial model, their value for the Gaussian model indicated that the best model explained a good portion of variability in the data (R^2^=0.75 ± 0.10) and had an intermediate correlation between observations (ICC=0.61, 89 % HDI = 0.57-0.66; between group variance = 0.51, 89 % HDI = 0.42-0.66).

**Fig. 3.**
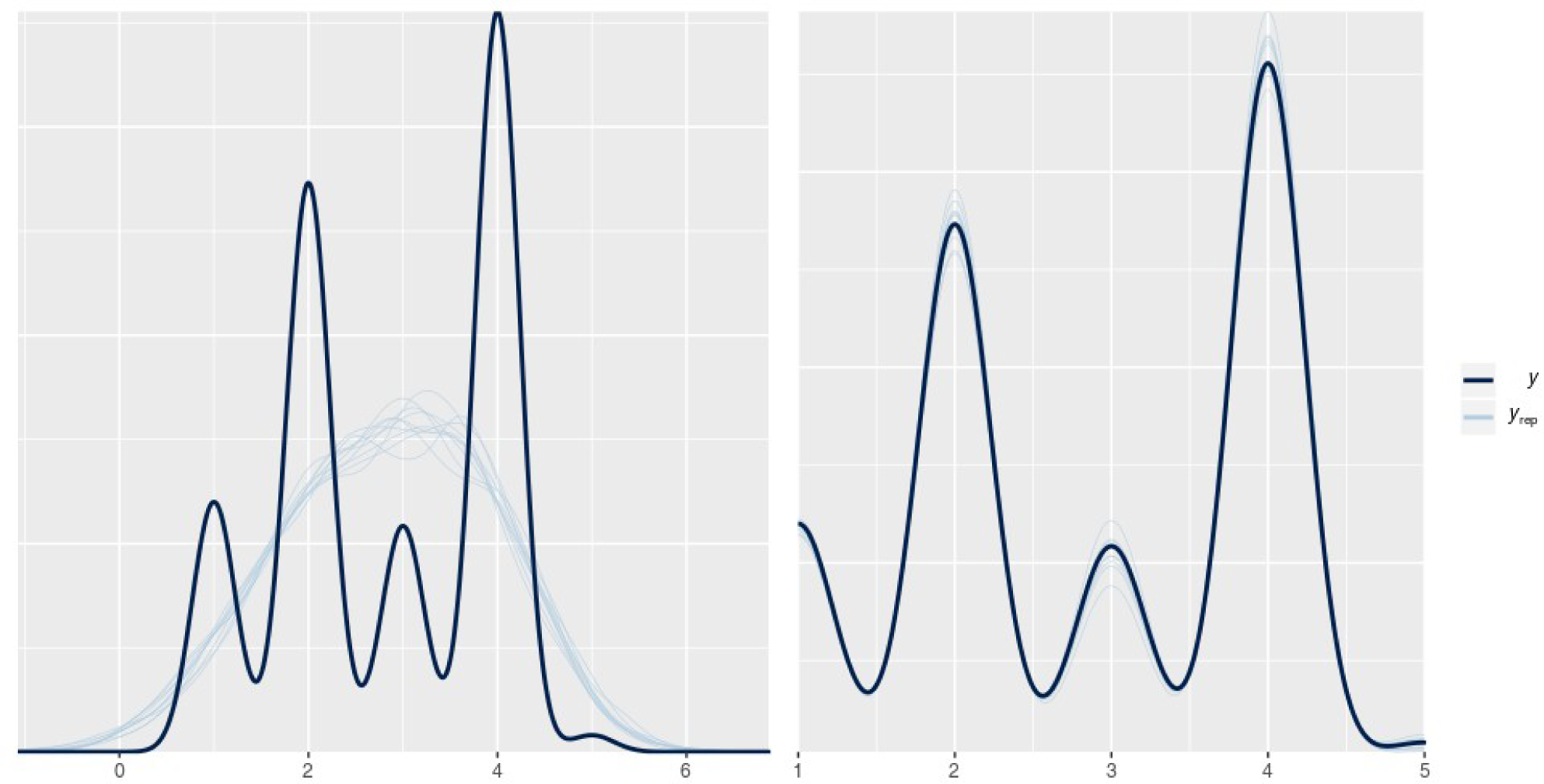
Distribution of the observed outcomes (dark blue line) with kernel density estimates of each replication from the posterior predictive distribution (light blue line). The model for ordinal answers (on the right), outperforms the Gaussian model (on the left).

Respondents believed that interactions between humans and ibexes were acceptable only when distances exceeded 25 meters. However, acceptability was lower the larger the group of hikers depicted in the picture: even when the group of people was at 50 meters from the ibex, at an acceptable distance, the situation was deemed to be unacceptable if there were more than 3 visitors. Very close observations of animals, with visitors at 5 meters, were always deemed unacceptable, regardless of group size (Fig. 4). Being a naturalistic photographer made respondents slightly more likely to rate a scenario as unacceptable (Table 1). The Potential for Conflict Index was similar, and relatively low, for all the combinations of distances and group size (Table 2).

**Fig. 4.**
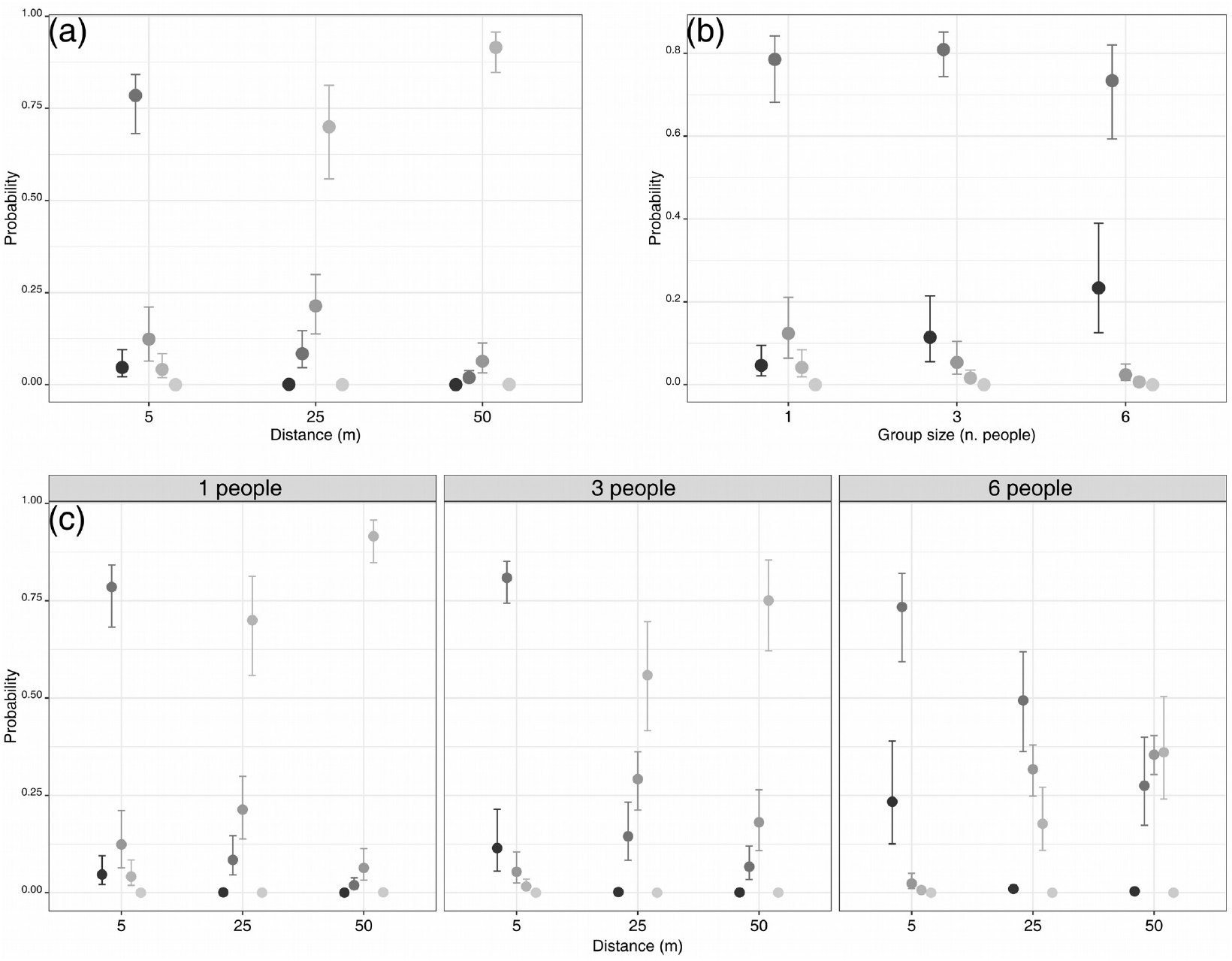
Acceptability of the various scenarios: marginal effect of distance (a), marginal effect of group size (b) and interaction between distance and group size (c).

**Table 1.** Output of the best candidate model: a Generalized Linear Mixed Model with a logarithmic link and a binomial structure of the error. The model has a random intercept and two random slopes for the covariates described in the vignettes: the distance between visitors and the ibex and the group size of visitors. Distance and group size are treated as Helmert contrasts.

| Group-level effects |  |  |  |  |  |  |
| --- | --- | --- | --- | --- | --- | --- |
| Variable | Estimate | Estimated Error | Lower 95% CI | Upper 95% CI | Eff. Sample | Rhat |
| sd (intercept) | 3.29 | 0.24 | 2.84 | 3.79 | 1910 | 1.00 |
| sd (Distance: 25 m) | 1.42 | 0.15 | 1.14 | 1.73 | 2573 | 1.00 |
| sd (Distance : 50 m) | 0.72 | 0.08 | 0.58 | 0.88 | 2697 | 1.00 |
| sd (Group size: 3 people) | 0.45 | 0.13 | 0.19 | 0.71 | 2408 | 1.00 |
| sd (Group size: 6 people) | 0.51 | 0.07 | 0.37 | 0.66 | 2414 | 1.00 |
| cor (Intercept, Distance: 25m) | -0.31 | 0.10 | -0.49 | -0.09 | 4819 | 1.00 |
| cor (Intercept, Distance: 50m) | -0.22 | 0.11 | -0.43 | 0.01 | 5392 | 1.00 |
| cor (Distance 25m, Distance: 50 m) | 0.88 | 0.06 | 0.75 | 0.97 | 2585 | 1.00 |
| cor (Intercept, Group size: 3 people) | 0.15 | 0.21 | -0.28 | 0.56 | 11492 | 1.00 |
| cor (Distance 25m, Group size: 3 people) | -0.19 | 0.22 | -0.60 | 0.24 | 6000 | 1.00 |
| cor (Distance 50m, Group size: 3 people) | -0.14 | 0.22 | -0.55 | 0.30 | 6357 | 1.00 |
| cor (Intercept, Group size: 6 people) | -0.14 | 0.14 | -0.41 | 0.12 | 7886 | 1.00 |
| cor (Distance 25m, Group size: 6 people) | -0.13 | 0.14 | -0.40 | 0.15 | 3772 | 1.00 |
| cor (Distance 50m, Group size: 5 people) | 0.01 | 0.15 | -0.28 | 0.29 | 3341 | 1.00 |
| cor (Group size: 25m, Group size: 50 m) | 0.72 | 0.16 | 0.32 | 0.94 | 1783 | 1.00 |
| Population-level effects |  |  |  |  |  |  |
| Variable | Estimate (log-odds) | Estimated Error | Lower 95% CI | Upper 95% CI | Eff. Sample | Rhat |
| Intercept [1] | -5.22 | 0.38 | -6.00 | -4.51 | 1568 | 1.00 |
| Intercept [2] | -0.58 | 0.32 | -1.22 | 0.04 | 1421 | 1.00 |
| Intercept [3] | 0.95 | 0.32 | 0.30 | 1.56 | 1394 | 1.00 |
| Intercept [4] | 10.43 | 0.68 | 9.16 | 11.82 | 2922 | 1.00 |
| Distance: 25 m | 1.96 | 0.15 | 1.67 | 2.27 | 3309 | 1.00 |
| Distance: 50 m | 1.03 | 0.08 | 0.87 | 1.20 | 3810 | 1.00 |
| Group size: 3 people | -0.48 | 0.09 | -0.65 | -0.30 | 9934 | 1.00 |
| Group size: 6 people | -0.64 | 0.06 | -0.77 | -0.52 | 5496 | 1.00 |
| Being a photographer | -0.33 | 0.48 | -1.29 | 0.60 | 1220 | 1.00 |
| Distance 25 m; Group size: 3 people | 0.09 | 0.09 | -0.10 | 0.27 | 18142 | 1.00 |

**Table 2.** Potential for conflict Index for each situation described in the vignettes

|  |  | Group size |  |  |
| --- | --- | --- | --- | --- |
|  |  | 1 people | 3 people | 6 people |
| Distance between visitors and the ibex | 5 meters | 0.32 | 0.25 | 0.23 |
|  | 25 meters | 0.26 | 0.23 | 0.22 |
|  | 50 meters | 0.21 | 0.26 | 0.24 |

## DISCUSSION

This research shows how a factorial survey based on manipulated digital pictures can help understanding human-wildlife interactions at protected areas and designing behavioral interventions aimed at fostering proper hiking behavior.

Our findings indicate that visitors hold encouraging perspectives about their potential interaction with alpine ibexes. Acceptable ibex observations are those occurring at more than 25 meters of distance and involving very small groups of visitors, with no more than 3 people. The most appropriate way to observe an alpine ibex is deemed to be one involving a single person who observes them at 50 meters of distance, or more. Moreover, hopefully, visitors deem totally unacceptable to approach ibexes at a close distance, for observing them. These results are corroborated by the relatively constant, and low, value of the Potential for Conflict Index, which reflects the lack of isolated groups of visitors with different perspectives about the acceptability of human-ibex interactions. The lack of groups with different beliefs will facilitate future conservation actions, as small groups of non-compliers are often critical for the effectiveness of management interventions.

When individual beliefs about a certain behavior are shared and cristallize around a common position, they pave the ground for the emergence of social norms, powerful informal institutions governing collective behavior (Bicchieri, 2016; Schultz, Nolan, Cialdini, Goldstein, & Griskevicius, 2018), which are a prerequisite for many nudging interventions. Leveraging on social norms can be an effective way for incentivize or inhibit the behaviors that are governed by these norms (e.g. through information campaigns and panels, Kinzig et al., 2013). We believe that this might be the case for our case study: overall, respondents shared common perspectives and a low disagreement on how wild ungulates like ibexes, should be observed. The Alpine ibex is a charismatic mammal, which is now common and easy to approach in many parts of the Alps. Therefore, our results could be useful for managers of areas where the interaction between ibex and tourists is perceived as a problem. Managers could enforce measurements aimed at promoting optimal distances for the observation of this species, especially at those hotspots where people are likely to approach animals. A potential intervention might include setting up panels at those hotspots. Panels can provide visitors with a normative statement, informing them that, according to our study, the majority of visitors of a national park believed that ibex should be observed by a maximum of 3 people from at least 25-30 meters. This statement, in its simplicity, could be a practical way to awake injunctive norms and to stimulate visitors to maintain a respectful distance from animals.

We believe that future studies should test and improve these panels. In this sense, it could be interesting to see whether adding some stimula, like human eyes (Nettle, Nott & Bateson, 2012), would improve their effectiveness over visitor behavior. Moreover, we believe that researchers should focus on how individual traits influence vignette evaluations. For reasons of time and field administration, in our study we did not explore how wildlife value orientations could influence the acceptability of human-ibex interactions. Value orientations are a major antecedents of human attitudes and behavior towards wildlife (Jacobs, Vaske, Teel & Manfredo, 2018) and assessing how they contribute to guide the evaluation of this interactions could be very useful to tailor communication panels or to optimize communication campaigns, targeting various groups of visitors (Miller, Freimund, Metcalf & Nickerson, 2018).

Finally, it is interesting to note that being a photographer or not was a predictor that was retained in our final model. On average, visitors who were naturalistic photographers were more severe in rating the acceptability of the various vignettes. This could happen because photographers are a specialized segments of recreationists, who can have a certain awareness about human disturbance to wildlife. Despite this awareness, photographers can locally become a source of stress for wild animals and behavioral interventions targeting them can be particularly important and beneficial. In this sense, it could be useful to identify role models or “positive deviants”, to enforce communication campaigns promoting a safe and respectful photographic activity.

## CONCLUSIONS

This study confirms that graphical factorial surveys based on digitally modified pictures can be an effective way to understand visitors’ acceptability of human-wildlife interactions at protected areas. While these type of study is relatively common in Northern America (Anderson, Manning, Valliere, & Hallo, 2010; Miller & Freimund, 2018), our research was arguably the first one in a European context and we believe that researchers could use it as a blueprint for studying human-wildlife interactions in Europe.

Our findings show that visitors of a large protected area in the Italian Alps have social norms governing how they approach large mammals when hiking. In plain terms, they have precise standards for defining whether a certain distance, or a certain group size, is appropriate for observing wildlife or not. The existence of these standards paves the ground for many behavioral interventions, like panels with normative statements, aimed at improving outdoor recreation while reducing negative effects of human-wildlife interactions.

## Supporting information

Supplementary dataset for reproducible data analysis

Digitally modified pictures adopted in the study

Software code for reproducible data analysis

## ACKNOWLEDGMENT

We are grateful to the Gran Paradiso National Park authority for their logistic support to this research.

