## Supplementary figures and images for "Graphical factorial surveys reveal the acceptability of wildlife observation at protected areas"

### 5m_GroupAlone.jpg

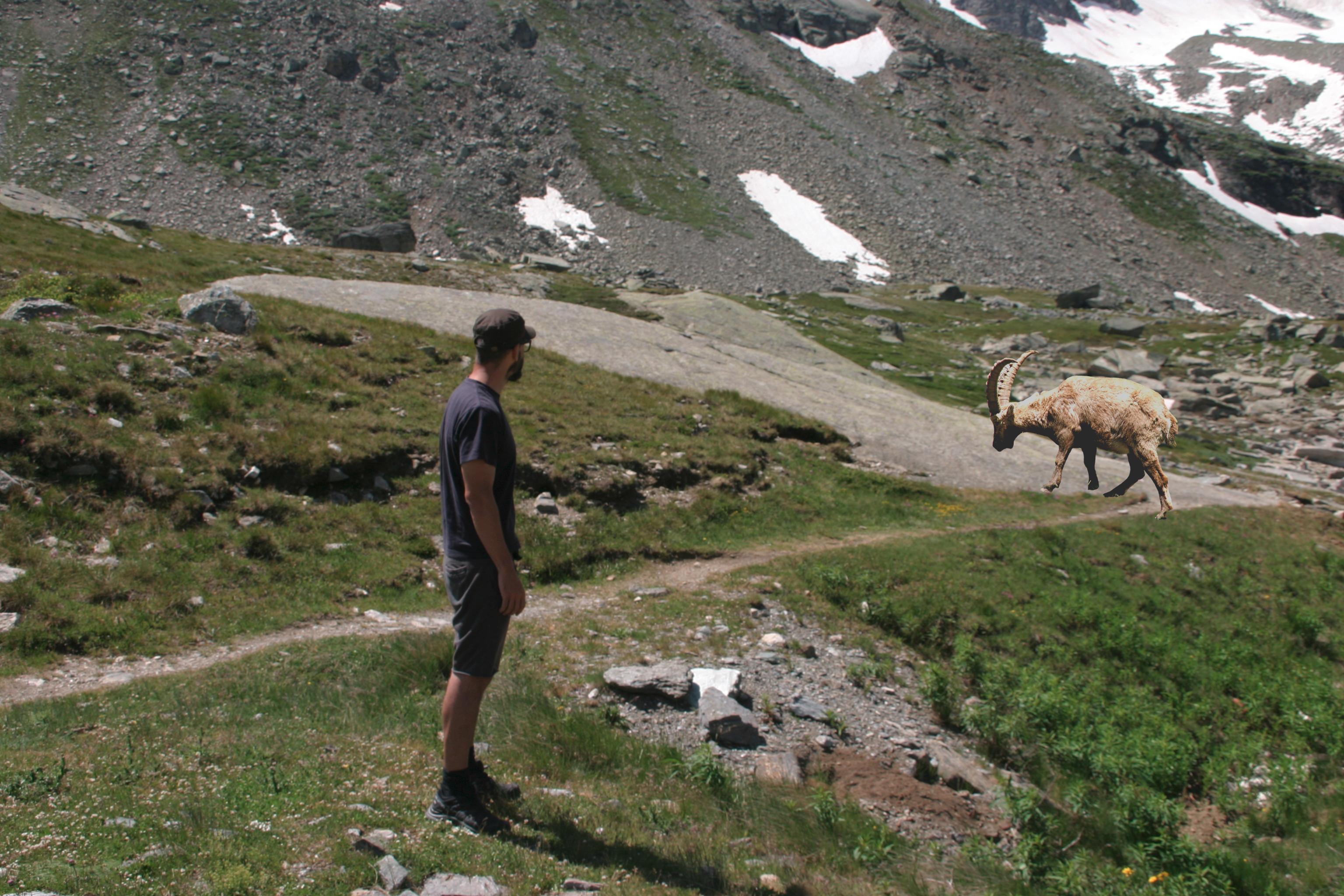

### 5m_GroupLarge.jpg

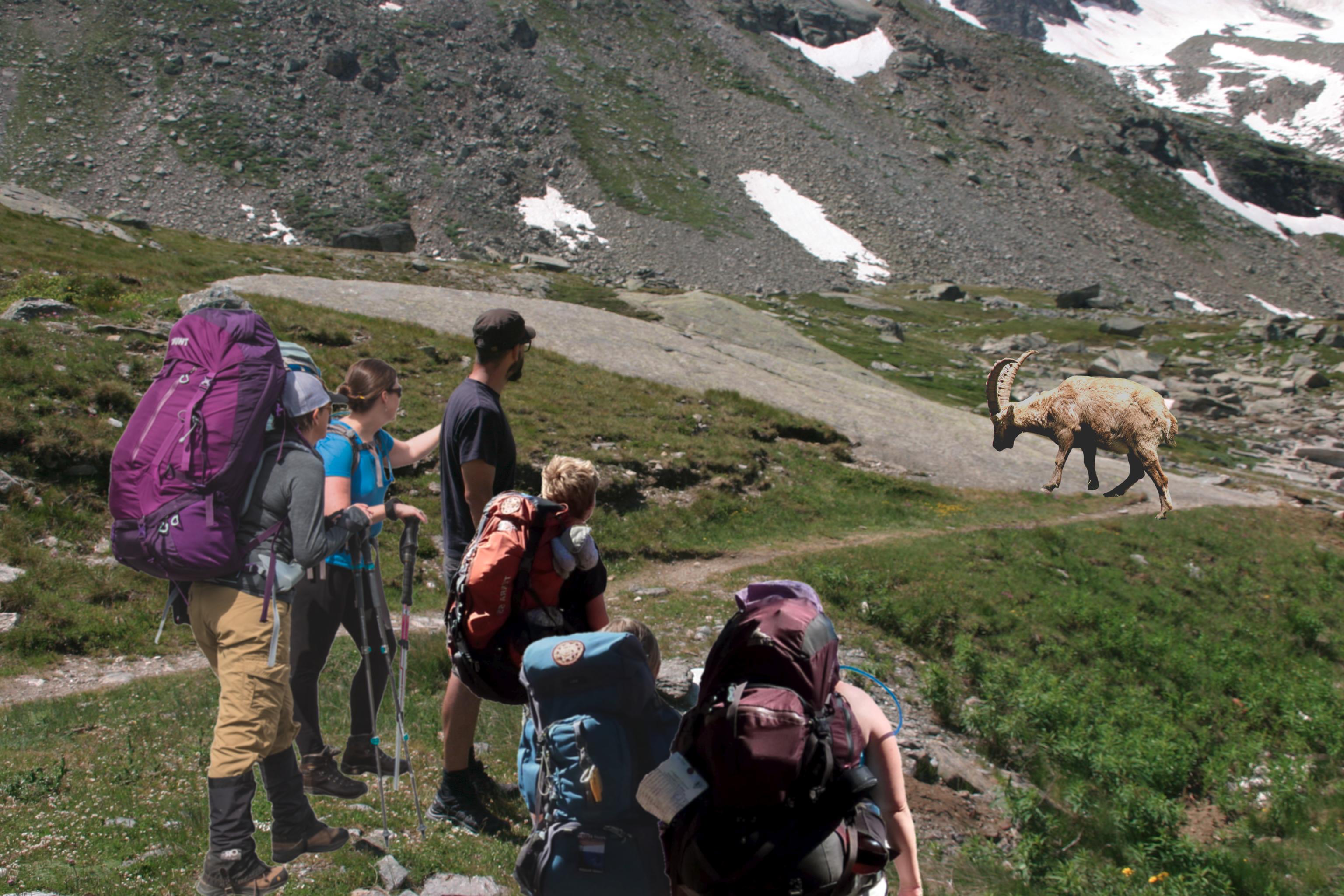

### 5m_GroupMed.jpg

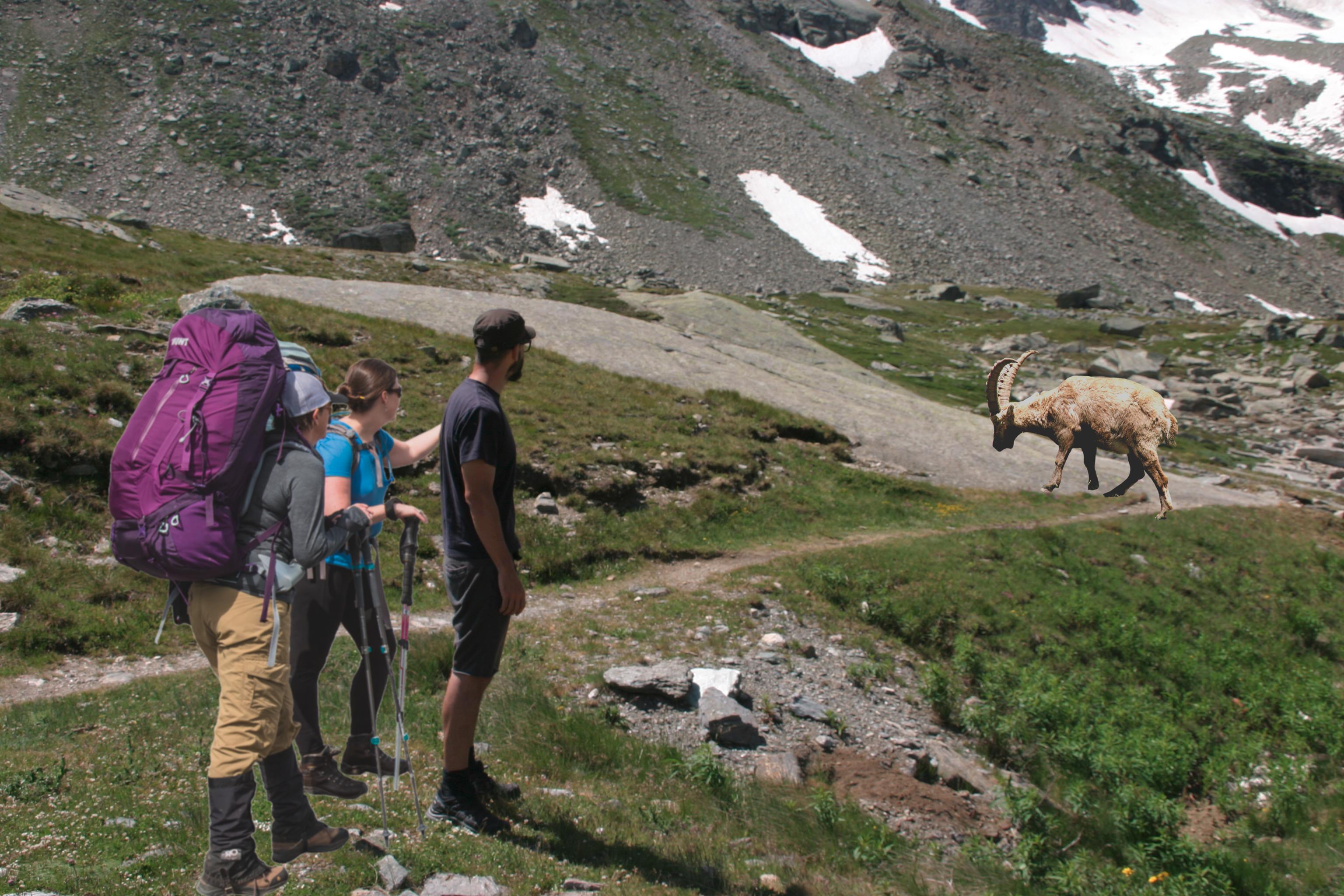

### 25m_GroupAlone.jpg

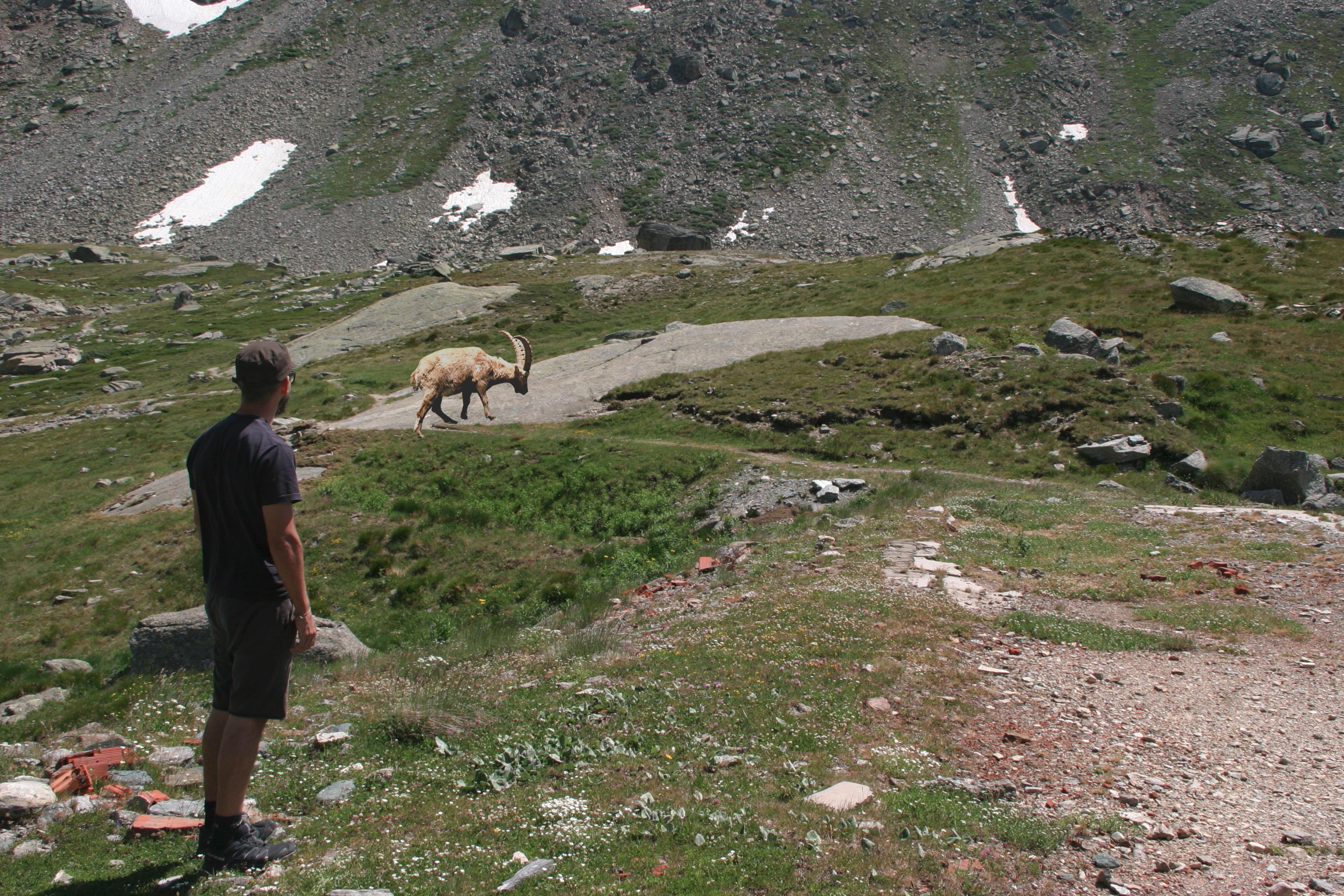

### 25m_GroupLarge.jpg

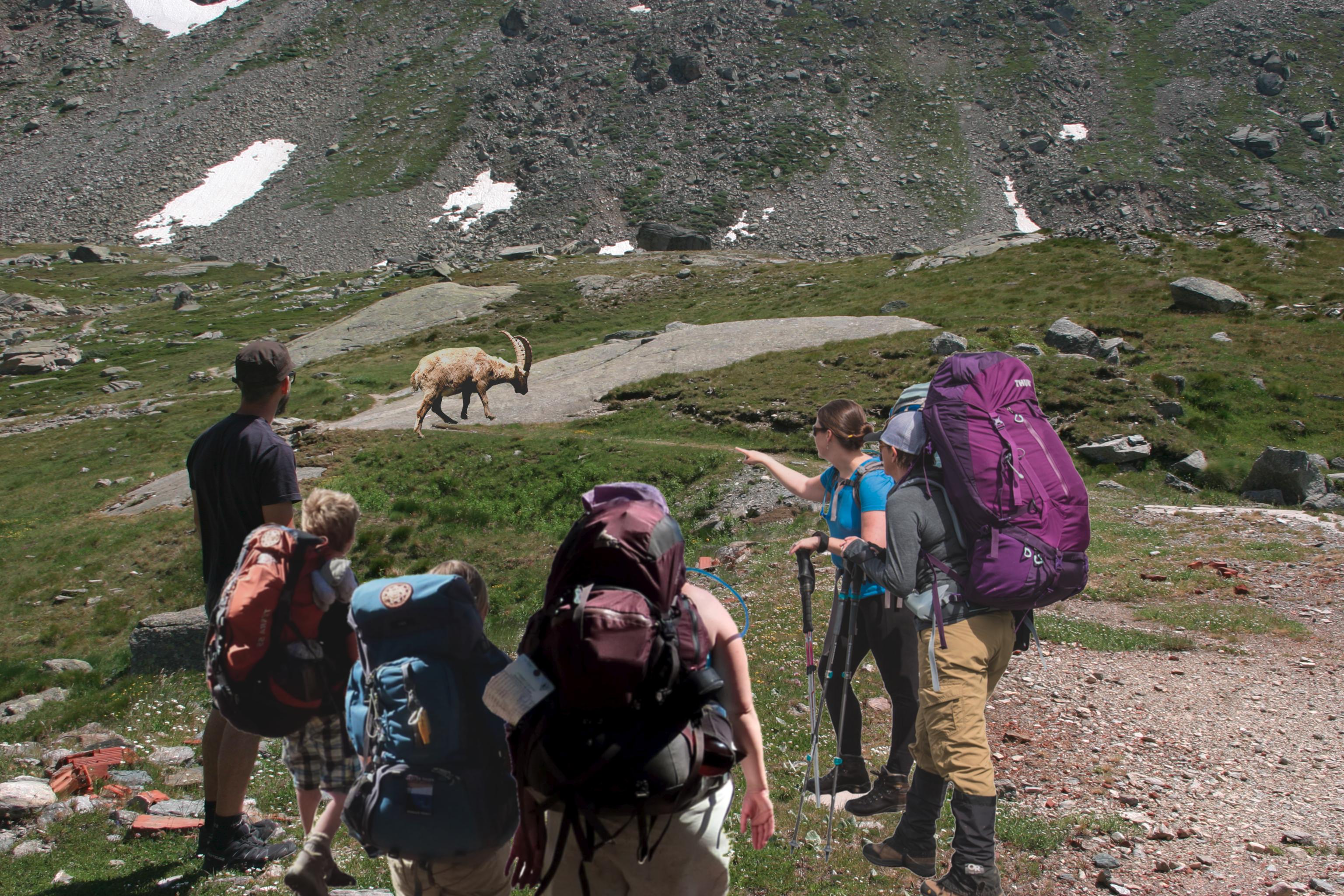

### 25m_GroupMed.jpg

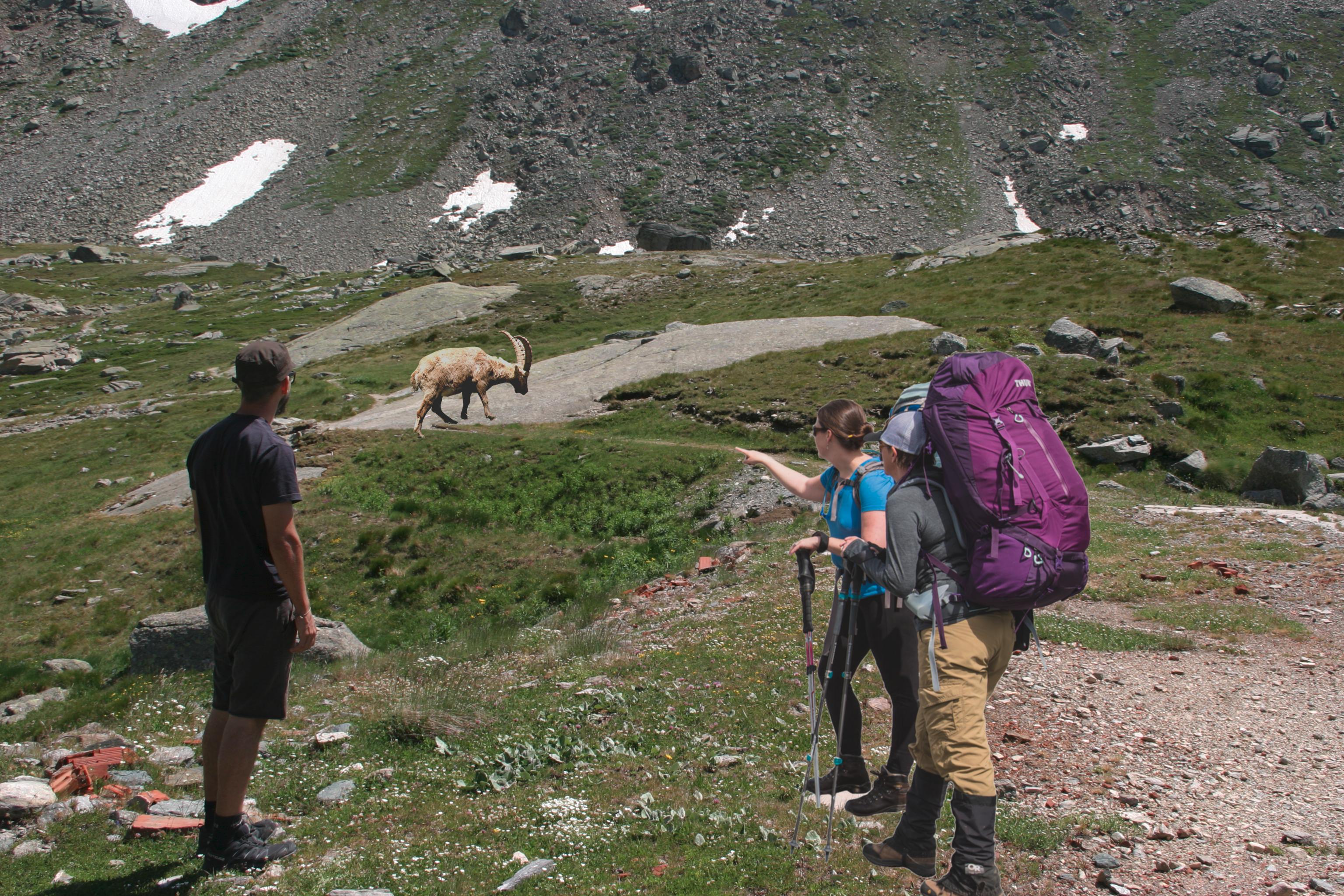

### 50m_GroupAlone.jpg

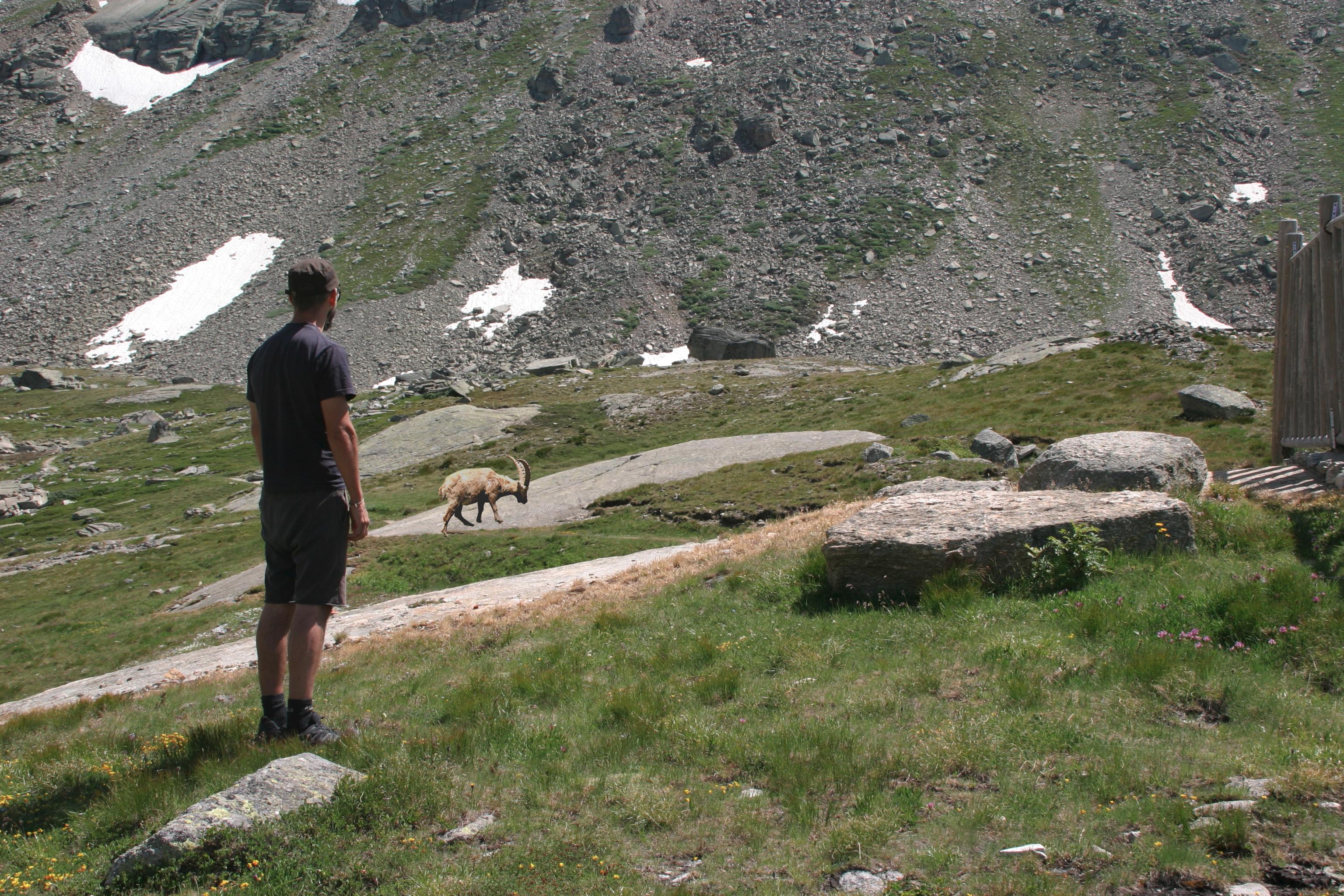

### 50m_GroupLarge.jpg

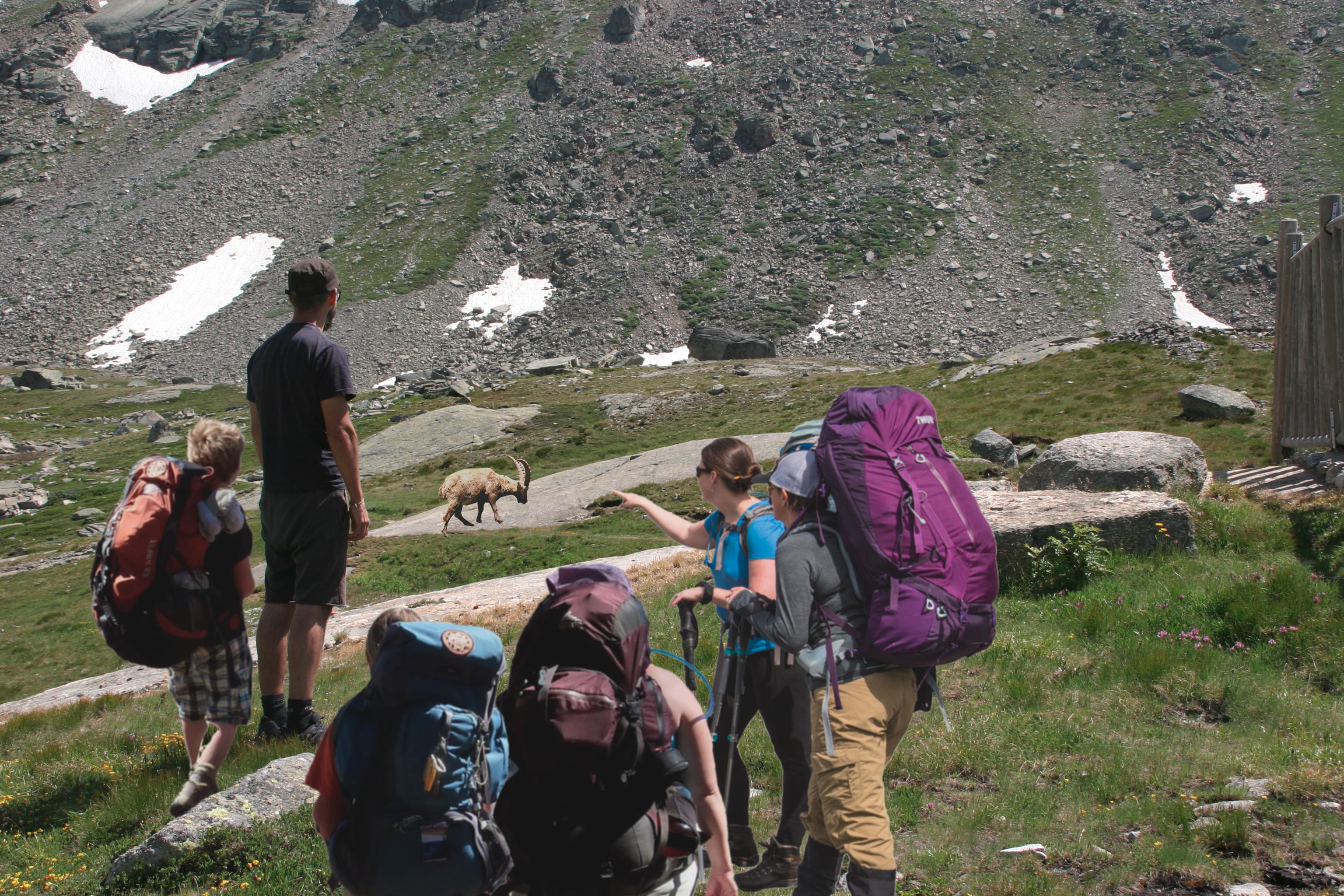

### 50m_GroupMed.jpg

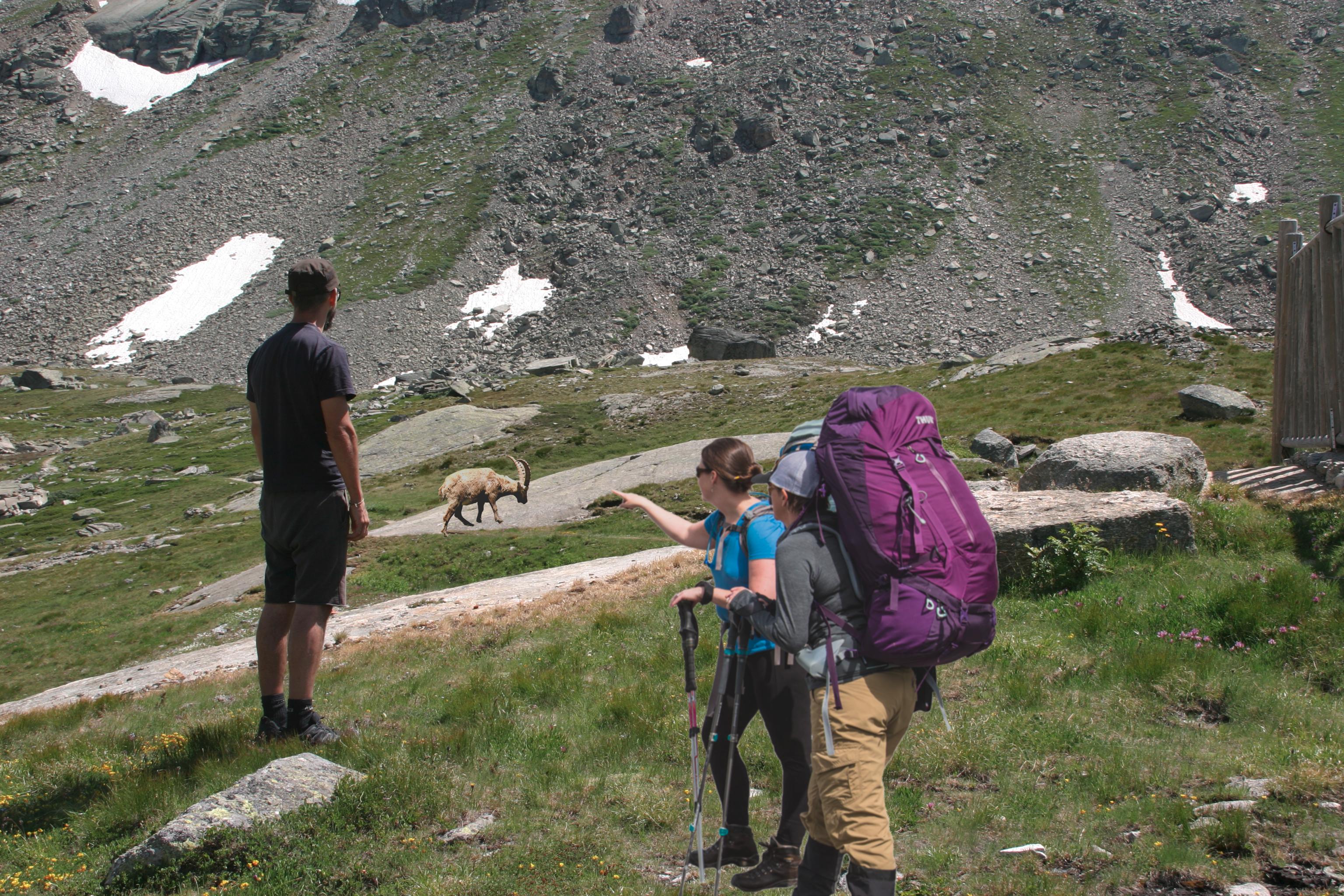
