## Supplementary material for "Graphical factorial surveys reveal the acceptability of wildlife observation at protected areas": Software code for reproducible data analysis: S2.R

```
#Loading packages
{
require(tidyverse)
require(devtools)
require(plyr)
require(reshape)
require(reshape2)
require(lubridate)
require(ggplot2)
require(brms)
#require(sjPlot)
#require(sjlabelled)
#require(sjmisc)
#require(sjstats)
require(jtools)
require(huxtable)
}

#################################################################
#Loading the dataset
#################################################################
{
dat <- read.csv(file="~/Downloads/granparadiso.csv")

#Removing respondents that did not know how to evaluate answers
head(dat)
names(dat)
levels(factor(dat$Respondent_ID)) #203 respondents in the original sample
levels(factor(dat[which(dat$Acceptability==1),"Respondent_ID"])) #0.24% people did not know how to evaluate
df <- dat[-which(dat$Acceptability==1),]
}

#################################################################
#Adjusting the dataset and setting contrasts
#################################################################
{
df$Group_size <- factor(df$Group_size)
df$Group_size <- plyr::revalue(df$Group_size, c("5"="6"))
contrasts(df$Group_size) <- "contr.helmert"

df$Distance <- factor(df$Distance)
table(df$Distance)
contrasts(df$Distance) <- "contr.helmert"

df$Acceptability <- factor(df$Acceptability)
names(df)
table(df$Acceptability)

prop.table(table(df$Hunter))
prop.table(table(df$Angler))
prop.table(table(df$Photo))
prop.table(table(df$EnvNGO))

df$Seen_ibex <- plyr::revalue(df$Seen_ibex, c("2 times"="Two times"))
df$Seen_ibex <- factor(df$Seen_ibex, levels=c("Never", "One time", "Two times", "Sometimes", "Often"))
prop.table(table(df$Seen_ibex))
contrasts(df$Seen_ibex) <- "contr.helmert"

levels(df$Frequency_visit)
df$Frequency_visit <- factor(df$Frequency_visit, c("First time", "Second time", "Once per year", "Multiple times per year",
                                                   "Multiple times per month", "Multiple times per week", "Once per week"))
prop.table(table(df$Frequency_visit))

hist(aggregate(Age ~ Respondent_ID, mean, data=df)$Age)
mean(dat$Age)
sd(dat$Age)
prop.table(table(df$Sex))

df[which(df$Education==""),"Education"] <- NA
df$Education <- factor(df$Education, levels=c("Secondary", "High", "Degree"))
prop.table(table(df$Education))
contrasts(df$Education) <- "contr.helmert"

table(df$Provenience)
prop.table(table(df$Peers))

table(df$Respondent_ID)
names(df)

#df$Photo <- factor(df$Photo)

table(df$Acceptability)
df$Acceptability <- ordered(df$Acceptability)

#Missing values
idx <- c("Acceptability", "Distance", "Group_size", "Photo", "Seen_ibex", "Respondent_ID")
foo <- df[,names(df)%in%idx]
fee <- foo[complete.cases(foo),]
nrow(fee)

tail(dat$Respondent_ID)
tail(dat$Respondent_ID)
dat$Date
table(dat$Respondent_ID)
table(dat$Place)
}

#################################################################
#Bayesian Hierarchical Modeling with STAN
#################################################################
{
modOrdered_Full <- brm(Acceptability ~ Distance + Group_size + Photo + Seen_ibex + Distance*Group_size*Photo +
              (Distance + Group_size|Respondent_ID), family="cumulative", data=fee, chains=4,
              iter=5000, cores = 6)
modOrdered_A <- brm(Acceptability ~ Distance + Group_size + Photo + Distance*Group_size*Photo +
                  (Distance + Group_size|Respondent_ID), family="cumulative", data=fee, chains=4,
                iter=5000, cores = 6)

modOrdered_B <- brm(Acceptability ~ Distance + Group_size + Photo + Distance*Group_size +
                  (Distance + Group_size|Respondent_ID), family="cumulative", data=fee, chains=4,
                iter=5000, cores = 6)

modOrdered_C <- brm(Acceptability ~ Distance + Group_size + Distance*Group_size +
                  (Distance + Group_size|Respondent_ID), family="cumulative", data=fee, chains=4,
                iter=5000, cores = 6)

save(modOrdered_Full, file="~/Documents/Jacopo/Research_Projects/Ongoing/Ibex_PNGP/Script/FullModel_Ordered.RData")
save(modOrdered_A, file="~/Documents/Jacopo/Research_Projects/Ongoing/Ibex_PNGP/Script/ModelA_Ordered.RData")
save(modOrdered_B, file="~/Documents/Jacopo/Research_Projects/Ongoing/Ibex_PNGP/Script/ModelB_Ordered.RData")
save(modOrdered_C, file="~/Documents/Jacopo/Research_Projects/Ongoing/Ibex_PNGP/Script/ModelC_Ordered.RData")


#Loading and selecting the best ordered GLMM
load(file="~/Documents/Jacopo/Research_Projects/Ongoing/Ibex_PNGP/Script/FullModel_Ordered.RData")
load(file="~/Documents/Jacopo/Research_Projects/Ongoing/Ibex_PNGP/Script/ModelA_Ordered.RData")
load(file="~/Documents/Jacopo/Research_Projects/Ongoing/Ibex_PNGP/Script/ModelB_Ordered.RData")
load(file="~/Documents/Jacopo/Research_Projects/Ongoing/Ibex_PNGP/Script/ModelC_Ordered.RData")

kappa1_Ordered <- loo(modOrdered_Full)
kappa2_Ordered <- loo(modOrdered_A)
kappa3_Ordered <- loo(modOrdered_B)
kappa4_Ordered <- loo(modOrdered_C)
leavoneoutsIC_Ordered <- c(kappa1_Ordered$looic, kappa2_Ordered$looic,
                           kappa3_Ordered$looic, kappa4_Ordered$looic)
wAIC_Ordered <- c(waic(modOrdered_Full)$waic, waic(modOrdered_A)$waic, waic(modOrdered_B)$waic,
                  waic(modOrdered_C)$waic)
bestmod_Ordered <- modOrdered_B
save(bestmod_Ordered, file="~/Documents/Jacopo/Research_Projects/Ongoing/Ibex_PNGP/Script/BestGLMM_Ordered.RData")

#Model with normal distribution
modGaussian_Full <- brm(as.numeric(Acceptability) ~ Distance + Group_size + Photo + Seen_ibex + Distance*Group_size*Photo +
                         (Distance + Group_size|Respondent_ID), family="Gaussian", data=fee, chains=4,
                       iter=5000, cores = 6)
modGaussian_A <- brm(as.numeric(Acceptability) ~ Distance + Group_size + Photo + Distance*Group_size*Photo +
                      (Distance + Group_size|Respondent_ID), family="Gaussian", data=fee, chains=4,
                    iter=5000, cores = 6)

modGaussian_B <- brm(as.numeric(Acceptability) ~ Distance + Group_size + Photo + Distance*Group_size +
                      (Distance + Group_size|Respondent_ID), family="Gaussian", data=fee, chains=4,
                    iter=5000, cores = 6)

modGaussian_C <- brm(as.numeric(Acceptability) ~ Distance + Group_size + Distance*Group_size +
                      (Distance + Group_size|Respondent_ID), family="Gaussian", data=fee, chains=4,
                    iter=5000, cores = 6)

save(modGaussian_Full, file="~/Documents/Jacopo/Research_Projects/Ongoing/Ibex_PNGP/Script/FullModel_Gaussian.RData")
save(modGaussian_A, file="~/Documents/Jacopo/Research_Projects/Ongoing/Ibex_PNGP/Script/ModelA_Gaussian.RData")
save(modGaussian_B, file="~/Documents/Jacopo/Research_Projects/Ongoing/Ibex_PNGP/Script/ModelB_Gaussian.RData")
save(modGaussian_C, file="~/Documents/Jacopo/Research_Projects/Ongoing/Ibex_PNGP/Script/ModelC_Gaussian.RData")

kappa1_Gaussian <- loo(modGaussian_Full)
kappa2_Gaussian <- loo(modGaussian_A)
kappa3_Gaussian <- loo(modGaussian_B)
kappa4_Gaussian <- loo(modGaussian_C)
leavoneoutsIC_Gaussian <- c(kappa1_Gaussian$looic, kappa2_Gaussian$looic,
                           kappa3_Gaussian$looic, kappa4_Gaussian$looic)
wAIC_Gaussian <- c(waic(modGaussian_Full)$waic, waic(modGaussian_A)$waic, waic(modGaussian_B)$waic,
                  waic(modGaussian_C)$waic)
bestmod_Gaussian <- modGaussian_B
save(bestmod_Gaussian, file="~/Documents/Jacopo/Research_Projects/Ongoing/Ibex_PNGP/Script/BestGLMM_Gaussian.RData")

#Comparing ordered logit and Gaussian GLMM
load(file="~/Documents/Jacopo/Research_Projects/Ongoing/Ibex_PNGP/Script/BestGLMM_Ordered.RData")
load(file="~/Documents/Jacopo/Research_Projects/Ongoing/Ibex_PNGP/Script/BestGLMM_Gaussian.RData")

summary(bestmod_Ordered)
summary(bestmod_Gaussian)

plot(bestmod_Ordered)

waic(bestmod_Ordered, bestmod_Gaussian)

pp_check(bestmod_Ordered)
pp_check(bestmod_Gaussian)

icc(bestmod_Gaussian, posterior=TRUE)
icc(bestmod, posterior=TRUE_Gaussian)

bayes_R2(bestmod_Gaussian)

#Plotting marginal effects
require(ggthemr)
require(RColorBrewer)
require(cowplot)
require(grid) # for unit()

ggthemr('earth')
p1 <- marginal_effects(bestmod_Ordered, categorical=TRUE, "Distance") 

pa <- plot(p1)[[1]] + ggplot2::labs(x = "Distance (m)", y = "Probability") + 
  scale_colour_grey() + theme_bw()+
  theme(axis.text.x = element_text(colour="black",size=12,angle=0,hjust=.5,vjust=.5,face="plain"),
        axis.text.y = element_text(colour="black",size=8,angle=0,hjust=1,vjust=0,face="plain"),  
        axis.title.x = element_text(colour="black",size=12,angle=0,hjust=.5,vjust=.5,face="plain"),
        axis.title.y = element_text(colour="black",size=12,angle=90,hjust=.5,vjust=.5,face="plain"),
        plot.margin = margin(t=33,  r=15, b=15, l=15)) + 
  theme(strip.text = element_text(colour = 'white', size=12)) + guides(fill=FALSE, color=FALSE)
pa

mylabels <- c("1", "3", "6")
p2 <- marginal_effects(bestmod_Ordered, categorical=TRUE, "Group_size") 
pb <- plot(p2)[[1]] + ggplot2::labs(x = "Group size (n. people)", y = "Probability") + 
  scale_colour_grey() + theme_bw()+ scale_x_discrete(labels=mylabels) +
  theme(axis.text.x = element_text(colour="black",size=12,angle=0,hjust=.5,vjust=.5,face="plain"),
        axis.text.y = element_text(colour="black",size=8,angle=0,hjust=1,vjust=0,face="plain"),  
        axis.title.x = element_text(colour="black",size=12,angle=0,hjust=.5,vjust=.5,face="plain"),
        axis.title.y = element_text(colour="black",size=12,angle=90,hjust=.5,vjust=.5,face="plain"),
        plot.margin = margin(t=33,  r=15, b=15, l=15)) + 
  theme(strip.text = element_text(colour = 'white', size=12)) + guides(fill=FALSE, color=FALSE)
pb

#mylabel <- c("1", "3", "5")
conditions <- data.frame(Group_size = c (1, 3, 5))
plotint <- marginal_effects(bestmod_Ordered, "Distance", 
                            conditions=conditions, categorical = TRUE)
pc <- plot(plotint)[[1]] + ggplot2::labs(x = "Distance (m)", y = "Probability") + 
  scale_colour_grey() + theme_bw()+
  theme(axis.text.x = element_text(colour="black",size=12,angle=0,hjust=.5,vjust=.5,face="plain"),
        axis.text.y = element_text(colour="black",size=8,angle=0,hjust=1,vjust=0,face="plain"),  
        axis.title.x = element_text(colour="black",size=12,angle=0,hjust=.5,vjust=0,face="plain"),
        axis.title.y = element_text(colour="black",size=12,angle=90,hjust=.5,vjust=.5,face="plain")) + 
  theme(strip.text = element_text(colour = 'white', size=12)) + guides(fill=FALSE, color=FALSE)
pc

#Multiple plot functions
require(grid)
pushViewport(viewport(layout = grid.layout(2, 2)))
vplayout <- function(x, y) viewport(layout.pos.row = x, layout.pos.col = y)
print(pc, vp = vplayout(1, 1:2))
print(pa, vp = vplayout(2, 1))
print(pb, vp = vplayout(2, 2))

ppord <- pp_check(bestmod_Ordered)
ppgaus <- pp_check(bestmod_Gaussian) + guides(col=FALSE)
pushViewport(viewport(layout = grid.layout(1, 2)))
vplayout <- function(x, y) viewport(layout.pos.row = x, layout.pos.col = y)
print(ppgaus, vp = vplayout(1, 1))
print(ppord, vp = vplayout(1, 2))


}

#################################################################
#Potential for Conflict Index
#################################################################
{
#Distance
#################################################################
{
table(fee$Distance, fee$Acceptability)

#Loading a frequency table
score <- c(-2:+2)
obs5m <- c(184, 243, 39, 106, 3)
obs25m <- c(31, 174, 99, 267, 5)
obs50m <- c(24, 127, 78, 335, 8)
dat <- data.frame(score,obs5m, obs25m, obs50m)
dat

#Specify distance and power (1 in this case)
d <- 1
p <- 1

#5 metres of distance
{
#Scale mean
Scale_mean5m <- sum(dat$score*dat$obs5m)/sum(obs5m)

#Total actual distance (TAD)
Tot_act_dist5m <- 
  (2*dat[which(dat$score==-2), "obs5m"]*dat[which(dat$score==1), "obs5m"]*((2+d-1)^p)) + 
  (2*dat[which(dat$score==-2), "obs5m"]*dat[which(dat$score==2), "obs5m"]*((3+d-1)^p)) +
  (2*dat[which(dat$score==-1), "obs5m"]*dat[which(dat$score==1), "obs5m"]*((1+d-1)^p)) +
  (2*dat[which(dat$score==-1), "obs5m"]*dat[which(dat$score==2), "obs5m"]*((2+d-1)^p))

Max_tot_dist5m <- if (sum(dat$obs5m) %% 2 == 0) {(((3+d-1)^p)*(sum(dat$obs5m))*((sum(dat$obs5m))/2))} else {
  (((3+d-1)^p)*(sum(dat$obs5m)+1)*((sum(dat$obs5m)-1)/2))}

#Observed PCI
PCI5m <- Tot_act_dist5m/Max_tot_dist5m
}

#25 metres of distance
{
  #Scale mean
  Scale_mean25m <- sum(dat$score*dat$obs25m)/sum(obs25m)
  
  #Total actual distance (TAD)
  Tot_act_dist25m <- 
    (2*dat[which(dat$score==-2), "obs25m"]*dat[which(dat$score==1), "obs25m"]*((2+d-1)^p)) + 
    (2*dat[which(dat$score==-2), "obs25m"]*dat[which(dat$score==2), "obs25m"]*((3+d-1)^p)) +
    (2*dat[which(dat$score==-1), "obs25m"]*dat[which(dat$score==1), "obs25m"]*((1+d-1)^p)) +
    (2*dat[which(dat$score==-1), "obs25m"]*dat[which(dat$score==2), "obs25m"]*((2+d-1)^p))
  
  Max_tot_dist25m <- if (sum(dat$obs25m) %% 2 == 0) {(((3+d-1)^p)*(sum(dat$obs25m))*((sum(dat$obs25m))/2))} else {
    (((3+d-1)^p)*(sum(dat$obs25m)+1)*((sum(dat$obs25m)-1)/2))}
  
  #Observed PCI
  PCI25m <- Tot_act_dist25m/Max_tot_dist25m
}

#50 metres of distance
{
  #Scale mean
  Scale_mean50m <- sum(dat$score*dat$obs50)/sum(obs50m)
  
  #Total actual distance (TAD)
  Tot_act_dist50m <- 
    (2*dat[which(dat$score==-2), "obs50m"]*dat[which(dat$score==1), "obs50m"]*((2+d-1)^p)) + 
    (2*dat[which(dat$score==-2), "obs50m"]*dat[which(dat$score==2), "obs50m"]*((3+d-1)^p)) +
    (2*dat[which(dat$score==-1), "obs50m"]*dat[which(dat$score==1), "obs50m"]*((1+d-1)^p)) +
    (2*dat[which(dat$score==-1), "obs50m"]*dat[which(dat$score==2), "obs50m"]*((2+d-1)^p))
  
  Max_tot_dist50m <- if (sum(dat$obs50m) %% 2 == 0) {(((3+d-1)^p)*(sum(dat$obs50m))*((sum(dat$obs50m))/2))} else {
    (((3+d-1)^p)*(sum(dat$obs50m)+1)*((sum(dat$obs50m)-1)/2))}
  
  #Observed PCI
  PCI50m <- Tot_act_dist50m/Max_tot_dist50m
}

PCI5m
PCI25m
PCI50m
}

#Group size
#################################################################
{
  table(fee$Group_size, fee$Acceptability)
  
  #Loading a frequency table
  score <- c(-2:+2)
  obs1p <- c(62, 149, 46, 307, 8)
  obs3p <- c(74, 169, 80, 243, 6)
  obs5p <- c(103, 226, 90, 158, 2)
  dat <- data.frame(score,obs1p, obs3p, obs5p)
  dat
  
  #Specify distance and power (1 in this case)
  d <- 1
  p <- 1
  
  #Single visitors
  {
    #Scale mean
    Scale_mean1p <- sum(dat$score*dat$obs1p)/sum(obs1p)
    
    #Total actual distance (TAD)
    Tot_act_dist1p <- 
      (2*dat[which(dat$score==-2), "obs1p"]*dat[which(dat$score==1), "obs1p"]*((2+d-1)^p)) + 
      (2*dat[which(dat$score==-2), "obs1p"]*dat[which(dat$score==2), "obs1p"]*((3+d-1)^p)) +
      (2*dat[which(dat$score==-1), "obs1p"]*dat[which(dat$score==1), "obs1p"]*((1+d-1)^p)) +
      (2*dat[which(dat$score==-1), "obs1p"]*dat[which(dat$score==2), "obs1p"]*((2+d-1)^p))
    
    Max_tot_dist1p <- if (sum(dat$obs1p) %% 2 == 0) {(((3+d-1)^p)*(sum(dat$obs1p))*((sum(dat$obs1p))/2))} else {
      (((3+d-1)^p)*(sum(dat$obs1p)+1)*((sum(dat$obs1p)-1)/2))}
    
    #Observed PCI
    PCI1p <- Tot_act_dist1p/Max_tot_dist1p
  }
  
  #Three visitors
  {
    #Scale mean
    Scale_mean3p <- sum(dat$score*dat$obs3p)/sum(obs3p)
    
    #Total actual distance (TAD)
    Tot_act_dist3p <- 
      (2*dat[which(dat$score==-2), "obs3p"]*dat[which(dat$score==1), "obs3p"]*((2+d-1)^p)) + 
      (2*dat[which(dat$score==-2), "obs3p"]*dat[which(dat$score==2), "obs3p"]*((3+d-1)^p)) +
      (2*dat[which(dat$score==-1), "obs3p"]*dat[which(dat$score==1), "obs3p"]*((1+d-1)^p)) +
      (2*dat[which(dat$score==-1), "obs3p"]*dat[which(dat$score==2), "obs3p"]*((2+d-1)^p))
    
    Max_tot_dist3p <- if (sum(dat$obs3p) %% 2 == 0) {(((3+d-1)^p)*(sum(dat$obs3p))*((sum(dat$obs3p))/2))} else {
      (((3+d-1)^p)*(sum(dat$obs3p)+1)*((sum(dat$obs3p)-1)/2))}
    
    #Observed PCI
    PCI3p <- Tot_act_dist3p/Max_tot_dist3p
  }
  
  #Five visitors
  {
    #Scale mean
    Scale_mean5p <- sum(dat$score*dat$obs5p)/sum(obs5p)
    
    #Total actual distance (TAD)
    Tot_act_dist5p <- 
      (2*dat[which(dat$score==-2), "obs5p"]*dat[which(dat$score==1), "obs5p"]*((2+d-1)^p)) + 
      (2*dat[which(dat$score==-2), "obs5p"]*dat[which(dat$score==2), "obs5p"]*((3+d-1)^p)) +
      (2*dat[which(dat$score==-1), "obs5p"]*dat[which(dat$score==1), "obs5p"]*((1+d-1)^p)) +
      (2*dat[which(dat$score==-1), "obs5p"]*dat[which(dat$score==2), "obs5p"]*((2+d-1)^p))
    
    Max_tot_dist5p <- if (sum(dat$obs5p) %% 2 == 0) {(((3+d-1)^p)*(sum(dat$obs5p))*((sum(dat$obs5p))/2))} else {
      (((3+d-1)^p)*(sum(dat$obs5p)+1)*((sum(dat$obs5p)-1)/2))}
    
    #Observed PCI
    PCI5p <- Tot_act_dist5p/Max_tot_dist5p
  }
  
  PCI1p
  PCI3p
  PCI5p
}

#Distance and group size
#################################################################
{
  
  xtabs(~ Acceptability + Group_size + Distance, data =fee)
  
  #Loading a frequency table
  score <- c(-2:+2)
  obs5m1p <- c(50, 76, 16, 45, 2)
  obs5m3p <- c(59, 84, 15, 33, 1)
  obs5m5p <- c(75, 83, 8, 28, 0)
  obs25m1p <- c(8, 43, 22, 118, 2)
  obs25m3p <- c(9, 46, 38, 97, 2)
  obs25m5p <- c(14, 85, 39, 52, 1)
  obs50m1p <- c(4, 30, 8, 144, 4)
  obs50m3p <- c(6, 39, 27, 113, 78)
  obs50m5p <- c(14, 58, 43, 78, 1)
  dat <- data.frame(score,obs5m1p, obs5m3p, obs5m5p,obs25m1p, obs25m3p, obs25m5p,
                    obs50m1p, obs50m3p, obs50m5p)
  dat
  
  #Specify distance and power (1 in this case)
  d <- 1
  p <- 1
  
  #Five metres, single visitors
  {
    #Scale mean
    Scale_mean5m1p <- sum(dat$score*dat$obs5m1p)/sum(obs5m1p)
    
    #Total actual distance (TAD)
    Tot_act_dist5m1p <- 
      (2*dat[which(dat$score==-2), "obs5m1p"]*dat[which(dat$score==1), "obs5m1p"]*((2+d-1)^p)) + 
      (2*dat[which(dat$score==-2), "obs5m1p"]*dat[which(dat$score==2), "obs5m1p"]*((3+d-1)^p)) +
      (2*dat[which(dat$score==-1), "obs5m1p"]*dat[which(dat$score==1), "obs5m1p"]*((1+d-1)^p)) +
      (2*dat[which(dat$score==-1), "obs5m1p"]*dat[which(dat$score==2), "obs5m1p"]*((2+d-1)^p))
    
    Max_tot_dist5m1p <- if (sum(dat$obs5m1p) %% 2 == 0) {(((3+d-1)^p)*(sum(dat$obs5m1p))*((sum(dat$obs5m1p))/2))}else {(((3+d-1)^p)*(sum(dat$obs5m1p)+1)*((sum(dat$obs5m1p)-1)/2))}
    
    #Observed PCI
    PCI5m1p <- Tot_act_dist5m1p/Max_tot_dist5m1p
  }

  #Five metres, three visitors
  {
    #Scale mean
    Scale_mean5m3p <- sum(dat$score*dat$obs5m3p)/sum(obs5m3p)
    
    #Total actual distance (TAD)
    Tot_act_dist5m3p <- 
      (2*dat[which(dat$score==-2), "obs5m3p"]*dat[which(dat$score==1), "obs5m3p"]*((2+d-1)^p)) + 
      (2*dat[which(dat$score==-2), "obs5m3p"]*dat[which(dat$score==2), "obs5m3p"]*((3+d-1)^p)) +
      (2*dat[which(dat$score==-1), "obs5m3p"]*dat[which(dat$score==1), "obs5m3p"]*((1+d-1)^p)) +
      (2*dat[which(dat$score==-1), "obs5m3p"]*dat[which(dat$score==2), "obs5m3p"]*((2+d-1)^p))
    
    Max_tot_dist5m3p <- if (sum(dat$obs5m3p) %% 2 == 0) {(((3+d-1)^p)*(sum(dat$obs5m3p))*((sum(dat$obs5m3p))/2))}else {(((3+d-1)^p)*(sum(dat$obs5m3p)+1)*((sum(dat$obs5m3p)-1)/2))}
    
    #Observed PCI
    PCI5m3p <- Tot_act_dist5m3p/Max_tot_dist5m3p
  }
  
  #Five metres, five visitors
  {
    #Scale mean
    Scale_mean5m5p <- sum(dat$score*dat$obs5m5p)/sum(obs5m5p)
    
    #Total actual distance (TAD)
    Tot_act_dist5m5p <- 
      (2*dat[which(dat$score==-2), "obs5m5p"]*dat[which(dat$score==1), "obs5m5p"]*((2+d-1)^p)) + 
      (2*dat[which(dat$score==-2), "obs5m5p"]*dat[which(dat$score==2), "obs5m5p"]*((3+d-1)^p)) +
      (2*dat[which(dat$score==-1), "obs5m5p"]*dat[which(dat$score==1), "obs5m5p"]*((1+d-1)^p)) +
      (2*dat[which(dat$score==-1), "obs5m5p"]*dat[which(dat$score==2), "obs5m5p"]*((2+d-1)^p))
    
    Max_tot_dist5m5p <- if (sum(dat$obs5m5p) %% 2 == 0) {(((3+d-1)^p)*(sum(dat$obs5m5p))*((sum(dat$obs5m5p))/2))}else {(((3+d-1)^p)*(sum(dat$obs5m5p)+1)*((sum(dat$obs5m5p)-1)/2))}
    
    #Observed PCI
    PCI5m5p <- Tot_act_dist5m5p/Max_tot_dist5m5p
  }
  
  #Twenty-five metres, single visitors
  {
    #Scale mean
    Scale_mean25m1p <- sum(dat$score*dat$obs25m1p)/sum(obs25m1p)
    
    #Total actual distance (TAD)
    Tot_act_dist25m1p <- 
      (2*dat[which(dat$score==-2), "obs25m1p"]*dat[which(dat$score==1), "obs25m1p"]*((2+d-1)^p)) + 
      (2*dat[which(dat$score==-2), "obs25m1p"]*dat[which(dat$score==2), "obs25m1p"]*((3+d-1)^p)) +
      (2*dat[which(dat$score==-1), "obs25m1p"]*dat[which(dat$score==1), "obs25m1p"]*((1+d-1)^p)) +
      (2*dat[which(dat$score==-1), "obs25m1p"]*dat[which(dat$score==2), "obs25m1p"]*((2+d-1)^p))
    
    Max_tot_dist25m1p <- if (sum(dat$obs25m1p) %% 2 == 0) {(((3+d-1)^p)*(sum(dat$obs25m1p))*((sum(dat$obs25m1p))/2))}else {(((3+d-1)^p)*(sum(dat$obs25m1p)+1)*((sum(dat$obs25m1p)-1)/2))}
    
    #Observed PCI
    PCI25m1p <- Tot_act_dist25m1p/Max_tot_dist25m1p
  }
  
  #Twenty-five metres, three visitors
  {
    #Scale mean
    Scale_mean25m3p <- sum(dat$score*dat$obs25m3p)/sum(obs25m3p)
    
    #Total actual distance (TAD)
    Tot_act_dist25m3p <- 
      (2*dat[which(dat$score==-2), "obs25m3p"]*dat[which(dat$score==1), "obs25m3p"]*((2+d-1)^p)) + 
      (2*dat[which(dat$score==-2), "obs25m3p"]*dat[which(dat$score==2), "obs25m3p"]*((3+d-1)^p)) +
      (2*dat[which(dat$score==-1), "obs25m3p"]*dat[which(dat$score==1), "obs25m3p"]*((1+d-1)^p)) +
      (2*dat[which(dat$score==-1), "obs25m3p"]*dat[which(dat$score==2), "obs25m3p"]*((2+d-1)^p))
    
    Max_tot_dist25m3p <- if (sum(dat$obs25m3p) %% 2 == 0) {(((3+d-1)^p)*(sum(dat$obs25m3p))*((sum(dat$obs25m3p))/2))}else {(((3+d-1)^p)*(sum(dat$obs25m3p)+1)*((sum(dat$obs25m3p)-1)/2))}
    
    #Observed PCI
    PCI25m3p <- Tot_act_dist25m3p/Max_tot_dist25m3p
  }
  
  #Twenty-five metres, five visitors
  {
    #Scale mean
    Scale_mean25m5p <- sum(dat$score*dat$obs25m5p)/sum(obs25m5p)
    
    #Total actual distance (TAD)
    Tot_act_dist25m5p <- 
      (2*dat[which(dat$score==-2), "obs25m5p"]*dat[which(dat$score==1), "obs25m5p"]*((2+d-1)^p)) + 
      (2*dat[which(dat$score==-2), "obs25m5p"]*dat[which(dat$score==2), "obs25m5p"]*((3+d-1)^p)) +
      (2*dat[which(dat$score==-1), "obs25m5p"]*dat[which(dat$score==1), "obs25m5p"]*((1+d-1)^p)) +
      (2*dat[which(dat$score==-1), "obs25m5p"]*dat[which(dat$score==2), "obs25m5p"]*((2+d-1)^p))
    
    Max_tot_dist25m5p <- if (sum(dat$obs25m5p) %% 2 == 0) {(((3+d-1)^p)*(sum(dat$obs25m5p))*((sum(dat$obs25m5p))/2))}else {(((3+d-1)^p)*(sum(dat$obs25m5p)+1)*((sum(dat$obs25m5p)-1)/2))}
    
    #Observed PCI
    PCI25m5p <- Tot_act_dist25m5p/Max_tot_dist25m5p
  }

  #Fifty metres, single visitors
  {
    #Scale mean
    Scale_mean50m1p <- sum(dat$score*dat$obs50m1p)/sum(obs50m1p)
    
    #Total actual distance (TAD)
    Tot_act_dist50m1p <- 
      (2*dat[which(dat$score==-2), "obs50m1p"]*dat[which(dat$score==1), "obs50m1p"]*((2+d-1)^p)) + 
      (2*dat[which(dat$score==-2), "obs50m1p"]*dat[which(dat$score==2), "obs50m1p"]*((3+d-1)^p)) +
      (2*dat[which(dat$score==-1), "obs50m1p"]*dat[which(dat$score==1), "obs50m1p"]*((1+d-1)^p)) +
      (2*dat[which(dat$score==-1), "obs50m1p"]*dat[which(dat$score==2), "obs50m1p"]*((2+d-1)^p))
    
    Max_tot_dist50m1p <- if (sum(dat$obs50m1p) %% 2 == 0) {(((3+d-1)^p)*(sum(dat$obs50m1p))*((sum(dat$obs50m1p))/2))}else {(((3+d-1)^p)*(sum(dat$obs50m1p)+1)*((sum(dat$obs50m1p)-1)/2))}
    
    #Observed PCI
    PCI50m1p <- Tot_act_dist50m1p/Max_tot_dist50m1p
  }
  
  #Fifty metres, three visitors
  {
    #Scale mean
    Scale_mean50m3p <- sum(dat$score*dat$obs50m3p)/sum(obs50m3p)
    
    #Total actual distance (TAD)
    Tot_act_dist50m3p <- 
      (2*dat[which(dat$score==-2), "obs50m3p"]*dat[which(dat$score==1), "obs50m3p"]*((2+d-1)^p)) + 
      (2*dat[which(dat$score==-2), "obs50m3p"]*dat[which(dat$score==2), "obs50m3p"]*((3+d-1)^p)) +
      (2*dat[which(dat$score==-1), "obs50m3p"]*dat[which(dat$score==1), "obs50m3p"]*((1+d-1)^p)) +
      (2*dat[which(dat$score==-1), "obs50m3p"]*dat[which(dat$score==2), "obs50m3p"]*((2+d-1)^p))
    
    Max_tot_dist50m3p <- if (sum(dat$obs50m3p) %% 2 == 0) {(((3+d-1)^p)*(sum(dat$obs50m3p))*((sum(dat$obs50m3p))/2))}else {(((3+d-1)^p)*(sum(dat$obs50m3p)+1)*((sum(dat$obs50m3p)-1)/2))}
    
    #Observed PCI
    PCI50m3p <- Tot_act_dist50m3p/Max_tot_dist50m3p
  }
  
  #Fifty metres, three visitors
  {
    #Scale mean
    Scale_mean50m5p <- sum(dat$score*dat$obs50m5p)/sum(obs50m5p)
    
    #Total actual distance (TAD)
    Tot_act_dist50m5p <- 
      (2*dat[which(dat$score==-2), "obs50m5p"]*dat[which(dat$score==1), "obs50m5p"]*((2+d-1)^p)) + 
      (2*dat[which(dat$score==-2), "obs50m5p"]*dat[which(dat$score==2), "obs50m5p"]*((3+d-1)^p)) +
      (2*dat[which(dat$score==-1), "obs50m5p"]*dat[which(dat$score==1), "obs50m5p"]*((1+d-1)^p)) +
      (2*dat[which(dat$score==-1), "obs50m5p"]*dat[which(dat$score==2), "obs50m5p"]*((2+d-1)^p))
    
    Max_tot_dist50m5p <- if (sum(dat$obs50m5p) %% 2 == 0) {(((3+d-1)^p)*(sum(dat$obs50m5p))*((sum(dat$obs50m5p))/2))}else {(((3+d-1)^p)*(sum(dat$obs50m5p)+1)*((sum(dat$obs50m5p)-1)/2))}
    
    #Observed PCI
    PCI50m5p <- Tot_act_dist50m5p/Max_tot_dist50m5p
  }
  
  pci <- c(PCI5m1p, PCI5m3p, PCI5m5p, PCI25m1p, PCI25m3p, PCI25m5p, PCI50m1p, PCI50m3p, PCI50m5p) 
  
}
}
```
